# Ecological trajectories and microbial network reorganization across caries-associated oral niches

**DOI:** 10.64898/2026.08.12.744380

**Authors:** Qingxiu Li, Lilan Zhao, Zhiwen Liu, Zhenjun Li

**Affiliations:** Institute of Infectious Diseases, Chinese Center for Disease Control and Prevention, 155 Changbai Road, Changping District, Beijing 102206, China; Department of Stomatology, The Second Xiangya Hospital, Central South University, Changsha, China; Changsha Stomatological Hospital, Changsha 410000, China

**Keywords:** Dental caries, oral microbiome, ecological trajectories, microbial association networks, predictive modelling

## Abstract

Dental caries is a biofilm-mediated disease associated with ecological changes in the oral microbiome. How microbial community organization differs among healthy plaque, caries-associated plaque, and carious dentin remains incompletely defined. We used 16S rRNA gene sequencing to profile paired supragingival plaque and carious dentin samples from patients with caries, together with supragingival plaque from healthy controls. Caries-associated plaque showed higher diversity than healthy plaque, whereas diversity was lower in carious dentin. Ecological ordering placed the three sample types along a health–plaque–dentin continuum. Association-network analysis showed distinct network structures in caries-associated plaque and carious dentin, with the dentin network displaying greater density and lower modularity. By integrating differential-abundance and network-centrality results, we identified taxa associated with the dentin niche. A sparse logistic-regression model using three genera distinguished plaque from dentin in patient-grouped cross-validation (AUROC, 0.780; AUPRC, 0.718). These cross-sectional findings describe niche-associated microbiome organization in dental caries and provide candidate features for future validation in independent, clinically relevant cohorts.

## Introduction

Dental caries is among the most prevalent chronic diseases worldwide and imposes a substantial health and economic burden [1,2]. Although it is classically described as a sugar-driven, acid-mediated process of enamel demineralization, caries is also an ecological disease involving coordinated changes in the oral microbiome [3–5]. In health, supragingival plaque is commonly enriched in commensal taxa that contribute to community stability and pH homeostasis, whereas dentin lesions are enriched in acidogenic and aciduric taxa adapted to low-pH, anaerobic conditions [6–9].

High-throughput sequencing has documented compositional differences between healthy and caries-associated communities, including enrichment of taxa such as Scardovia and Lactobacillus in lesions [10,11]. However, most studies compare health and disease as static endpoints. This approach provides limited information about community organization in plaque from people with caries or about the relationship between plaque and the underlying dentin lesion. In addition, many analyses focus on taxon abundance rather than on changes in microbial association structure [12–15].

Here, we used paired sampling to compare supragingival plaque and carious dentin from the same patients, with healthy supragingival plaque as a reference group. We integrated compositional profiling, diversity analysis, ecological ordering, and microbial association-network analysis to describe niche-associated community patterns. We also evaluated whether a sparse, internally validated model could distinguish caries-associated plaque from carious dentin. Because the study is cross-sectional, all ordering and modelling results are interpreted as associations across sampled niches rather than as direct evidence of temporal progression, causal interactions, or clinical utility.

## Materials and Methods

### Study design and participant recruitment

This study used a paired sampling design to characterize niche-associated microbial communities in dental caries. Supragingival plaque and carious dentin were collected from the same individuals diagnosed with dental caries, and supragingival plaque was collected from healthy controls. Participants were recruited from the Department of Stomatology, The Second Xiangya Hospital, Central South University. Caries diagnoses were independently confirmed by two calibrated clinicians according to the World Health Organization Oral Health Surveys (5th edition). Inclusion criteria included at least 20 natural teeth, no systemic disease, and no antibiotic use within the preceding month. Individuals with a history of smoking, pregnancy, orthodontic treatment, or recent antimicrobial exposure were excluded.

Participants were classified as adolescents (12–18 years) or adults (≥19 years) for covariate analyses; age was not a primary stratification factor. The final dataset comprised 67 samples: supragingival plaque from seven healthy controls (H_sp), and paired supragingival plaque (D_sp) and carious dentin (D_ct) samples from 30 patients with caries. Supragingival plaque was collected using sterile Gracey curettes, and carious dentin was collected using sterile spoon excavators. Samples were placed on ice immediately after collection, stored at −20 °C, and transferred to −80 °C within 2 h.

Written informed consent was obtained from all participants prior to sampling. The study protocol was approved by the Ethics Committee of Xiangya Second Hospital, Central South University (Approval No. 2021-Ethics-Review-Clinical-021). Participant characteristics and sample distribution are summarized in Table 1.

**Table 1.** Participant and sample characteristics.

| Characteristic | H_sp | D_sp | D_ct |
| --- | --- | --- | --- |
| Number of samples | 7 | 30 | 30 |
| Number of participants | 7 | 30 | 30 |
| Sex (male/female) | 3/4 | 11/19 | 11/19 |
| Age, median (range), years | 35 (19–67) | 22 (12–48) | 22 (12–48) |
| Age group (adolescent/adult) | 0/7 | 15/15 | 15/15 |
| DMFT score, median | 0 | ≥3 | ≥3 |
| Sample type | Supragingival plaque | Supragingival plaque | Carious dentin |

H\_sp, healthy supragingival plaque; D\_sp, caries-associated supragingival plaque; D\_ct, carious dentin. D\_sp and D\_ct were paired samples from the same 30 participants with caries. DMFT, decayed, missing, and filled teeth.

### DNA extraction and amplicon sequencing

Genomic DNA was extracted from microbial pellets by SDS–proteinase K lysis, phenol–chloroform extraction, and ethanol precipitation. DNA integrity was assessed by agarose gel electrophoresis, and DNA concentration was quantified with a Qubit 3.0 fluorometer (Thermo Fisher Scientific). The V3– V4 region of the bacterial 16S rRNA gene was amplified with primers 341F and 806R using Phusion High-Fidelity PCR Master Mix (New England Biolabs). PCR comprised 30 cycles with a 56 °C annealing step. All samples were amplified in one batch to minimize batch effects. Amplicons were purified with AMPure XP beads, quantified using an Agilent 2100 Bioanalyzer, pooled at equimolar concentrations, and sequenced on an Illumina HiSeq 2500 platform (2 × 300 bp, paired-end). Negative controls were included during extraction and amplification and yielded negligible reads.

### Sequence processing and taxonomic assignment

Raw reads were processed in R (v4.3.2) using DADA2 (v1.28.0) [16]. Reads were quality filtered (Phred score ≥30), truncated to 280 bp (forward) and 220 bp (reverse), dereplicated, error-corrected, and merged with a minimum overlap of 20 bp. Chimeras were removed with the consensus method. Amplicon sequence variants (ASVs) were assigned taxonomically against SILVA release 138.1 using a naïve Bayesian classifier with a minimum bootstrap confidence of 50. To reduce sparsity in downstream analyses, ASVs with <0.1% relative abundance or detected in <5% of samples were excluded. On average, approximately 49,000 high-quality reads per sample were retained, yielding 3,752 ASVs. Genus-level abundances were used for community-composition and predictive-modelling analyses.

### Diversity analysis and phylogenetic reconstruction

Alpha diversity was quantified by Shannon diversity and Chao1 richness in phyloseq (v1.44.0) after rarefaction to 20,000 reads per sample [17]. The depth was selected to balance sequencing depth and sample retention; sensitivity analyses using non-rarefied data gave consistent diversity patterns. Pairwise and multi-group differences were evaluated with Wilcoxon rank-sum and Kruskal–Wallis tests, respectively, with Benjamini–Hochberg false-discovery-rate (FDR) correction [18–21]. Beta diversity was calculated with Bray–Curtis dissimilarities and visualized by principal coordinates analysis (PCoA). Group differences were tested by PERMANOVA (adonis, vegan v2.6-6; 999 permutations), and homogeneity of multivariate dispersion was assessed with betadisper. Representative ASV sequences were aligned using MAFFT (v7.520), and maximum-likelihood phylogenetic trees were constructed with IQ-TREE (v2.2.5; 1,000 bootstrap replicates) [22,23]. Trees were visualized with GraPhlAn (v1.1.4) and ggtree (v3.8.2) [24,25].

### Differential taxon analysis and ecological trajectory inference

Differentially abundant taxa were identified with DESeq2 (v1.42.0) using a paired design that included PatientID as a blocking factor and sampling location (H_sp, D_sp, and D_ct) as the primary fixed effect [26]. LEfSe was used as a complementary analysis (α = 0.05; LDA score >2.0) [27]. Ecological ordering was explored with monocle3 (v1.3.4) and principal-curve fitting using Bray–Curtis dissimilarities derived from ASV profiles [28–30]. H_sp samples were used as the reference endpoint. These analyses were used to visualize cross-sectional ordering across niches and were not interpreted as temporal reconstruction. Generalized additive models were used to describe taxa associated with positions along the H_sp–D_sp–D_ct ordering [5,31].

### Network reconstruction and decision-support modeling

Microbial association networks were constructed separately for D_sp and D_ct from ASV-level abundance profiles; node labels report genus-level taxonomy. SparCC was used to account for compositionality [32], and associations were retained only when they were also supported by Spearman correlation (|ρ| >0.6; FDR-adjusted P <0.05) [34]. Network topology metrics, including degree, betweenness centrality, and modularity, were calculated with igraph (v1.4.3) and visualized with ggraph (v2.1.0). Differential associations between D_sp and D_ct were assessed using NetCoMi (v1.0.5) under a permutation-based framework [35]. These correlation networks describe statistical associations and do not establish direct ecological interactions.

For predictive modelling, centered log-ratio (CLR)-transformed genus abundances were used to fit a sparse logistic-regression model with glmnet (v4.1-7) [36]. Regularization was selected using the λ_1se criterion to favor parsimony. Performance was evaluated using patient-grouped cross-validation to prevent leakage between paired samples. Discrimination was summarized by the area under the receiver-operating-characteristic curve (AUROC) and the area under the precision–recall curve (AUPRC); calibration was assessed by comparing predicted probabilities with observed class frequencies. All analyses were conducted in R (v4.3.2), with FDR <0.05 considered statistically significant. Figures were generated using ggplot2, ggraph, and ComplexHeatmap [37,38].

## Results

### 1. Community structure across oral niches

We sequenced the V3–V4 region of the 16S rRNA gene from 67 samples, yielding approximately 1.1 million high-quality reads and 6,025 ASVs. After prevalence and abundance filtering, 1,050 ASVs were retained for downstream analyses. Across all samples, the oral microbiota was dominated by Bacteroidetes, Actinobacteria, Fusobacteria, and Firmicutes, with additional contributions from Proteobacteria and Saccharibacteria. Prevotella, Leptotrichia, Capnocytophaga, Fusobacterium, Corynebacterium, Actinomyces, Porphyromonas, Veillonella, and Saccharibacteria (TM7) lineages were detected across oral niches (Fig. 1A).

**Figure 1.**
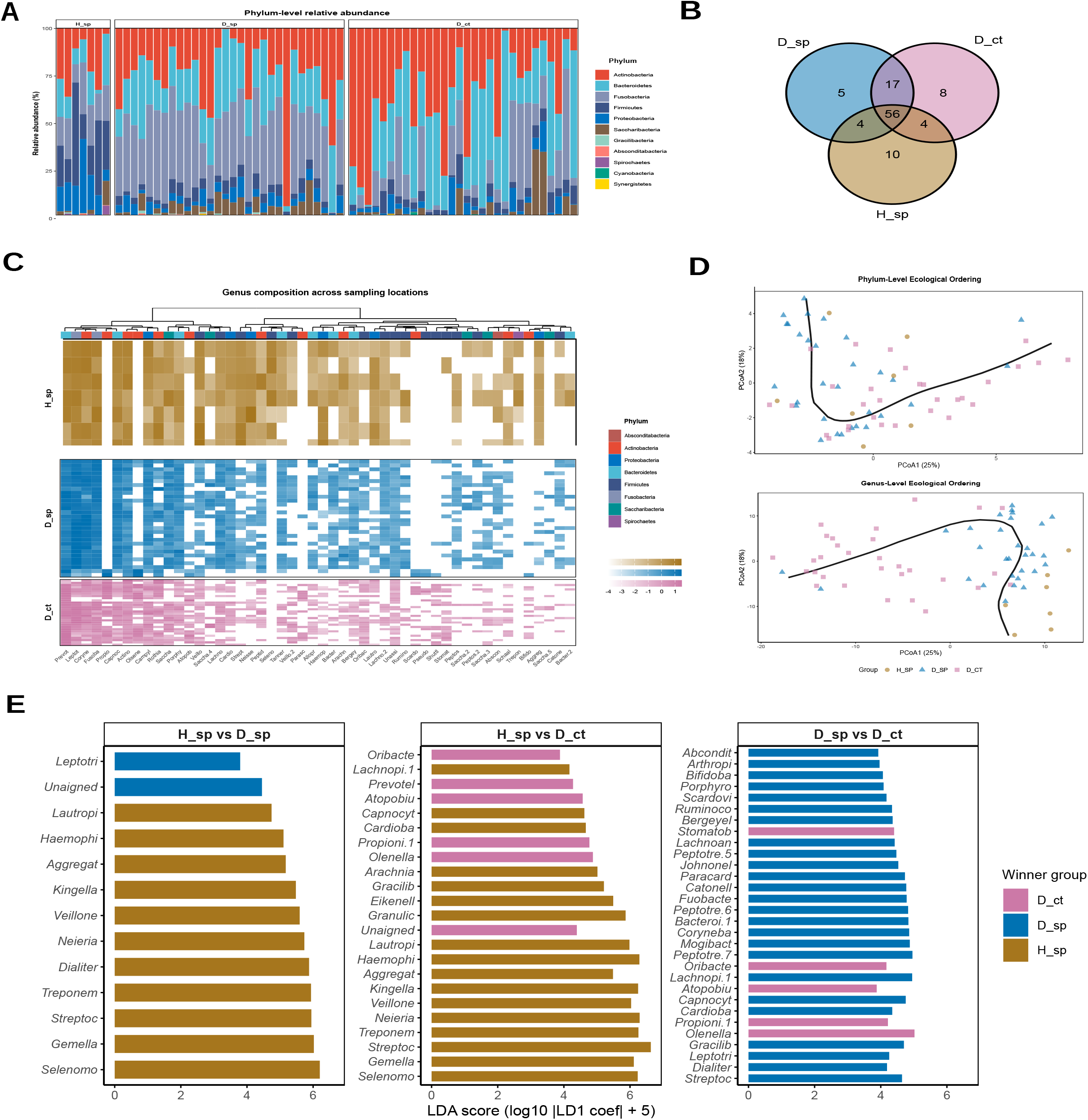
Community structure across oral niches. (A) Genus-level phylogenetic reconstruction, with branches coloured by phylum. (B) Shannon diversity by disease status, sampling location, sex, dentition status, and age group. (C) Principal-coordinate analysis of Bray–Curtis dissimilarities; the disease-status and sampling-location comparisons are shown alongside host-associated strata. H_sp, healthy supragingival plaque; D_sp, caries-associated supragingival plaque; D_ct, carious dentin.

### 2. Diversity patterns across plaque and dentin

Shannon diversity increased from healthy supragingival plaque (H_sp; median, 3.854) to caries-associated supragingival plaque (D_sp; median, 5.459; Wilcoxon test, FDR = 3.88 × 10^−4^) and was lower in carious dentin (D_ct; median, 5.072; FDR = 3.88 × 10^−4^) (Fig. 1B). Chao1 medians were 141, 441, and 316 for H_sp, D_sp, and D_ct, respectively. These cross-sectional data show non-linear diversity differences among the three sampled niches.

Bray–Curtis ordination showed separation by sampling location, which explained 11.5% of community variation (PERMANOVA, P = 0.001; Fig. 1C). Gender (R^2^ = 0.002, P = 0.064), dentition status (R^2^ = 0.018, P = 0.178), and age group (R^2^ = 0.017, P = 0.282) each explained <2% of variation. Within-group dispersion also differed (betadisper, P = 0.006); therefore, the PERMANOVA result should be interpreted as evidence of overall compositional separation rather than a difference in group centroids alone.

### 3. Taxonomic restructuring across oral niches

Pronounced compositional differences were observed across oral niches at both the phylum and genus levels. Healthy supragingival plaque (H_sp) was enriched in aerobic and facultative commensals, including *Neisseria, Haemophilus*, and *Corynebacterium*, whereas carious dentin (D_ct) was dominated by acidogenic and aciduric taxa such as *Scardovia, Lactobacillus*, and *Prevotella* (Fig. 2A). In contrast, caries-associated supragingival plaque (D_sp) displayed an intermediate taxonomic profile, characterized by enrichment of facultative anaerobes such as *Leptotrichia*, consistent with its position between health-associated and lesion-associated communities.

**Figure 2.**
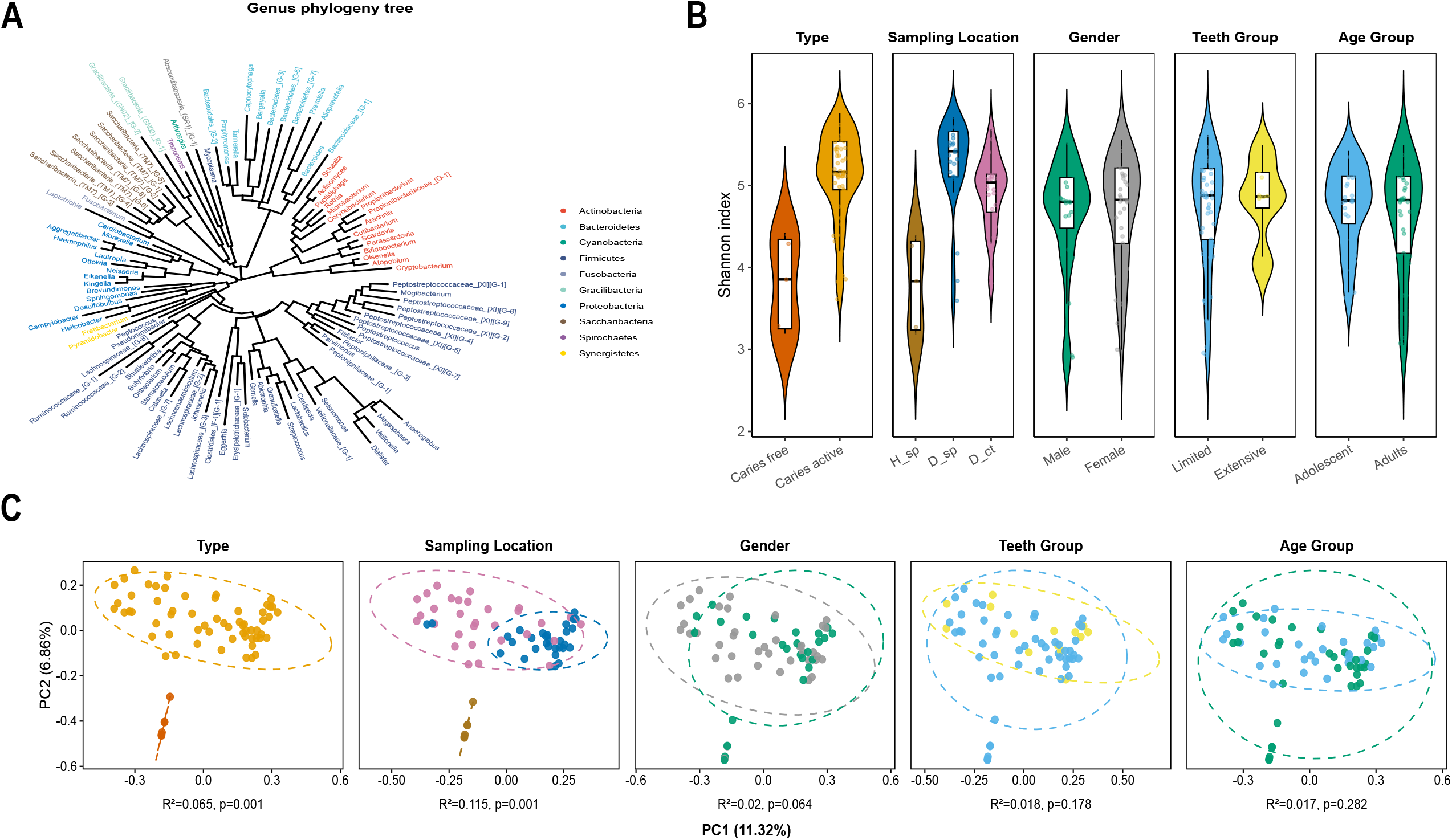
Taxonomic reorganization and ecological ordering across healthy plaque, caries-associated plaque, and carious dentin. (A) Phylum-level relative abundance across samples. (B) Venn diagram of genera shared among H_sp, D_sp, and D_ct. (C) Genus-level composition heatmap. (D) Principal-coordinate ecological ordering based on phylum- and genus-level Bray–Curtis dissimilarities; it is not a temporal reconstruction. (E) Pairwise LEfSe analysis of discriminative genera. H_sp, healthy supragingival plaque; D_sp, caries-associated supragingival plaque; D_ct, carious dentin.

Core–accessory analysis identified 56 genera shared across all niches, alongside niche-specific taxa, including ten unique to H_sp, one unique to D_sp, and eight unique to D_ct (Fig. 2B). Fermentative anaerobes were largely restricted to dentin lesions, reflecting adaptation to acidic and oxygen-depleted microenvironments. Genus-level heatmaps further supported these patterns, with H_sp and D_sp clustering closely, whereas D_ct communities exhibited marked restructuring, characterized by expansion of lesion-associated taxa (e.g., *Scardovia, Olsenella*) and contraction of health-associated commensals (e.g., *Neisseria, Haemophilus*) (Fig. 2C). Collectively, these results demonstrate stepwise taxonomic reorganization across oral niches.

### 4. Ecological ordering across plaque and dentin

Ecological ordering aligned H_sp, D_sp, and D_ct samples along a continuous axis, with D_sp occupying an intermediate position (Fig. 2D). Paired differential-abundance analysis identified taxa that differed across niches. Scardovia was enriched in dentin lesions, whereas Streptococcus showed a modest decline; Lactobacillus showed relatively stable abundance across the sampled niches. LEfSe identified 25 discriminatory genera (LDA >2.0; FDR <0.05; Fig. 2E). Neisseria, Haemophilus, and Corynebacterium were enriched in H_sp; Leptotrichia and Veillonella were enriched in D_sp; and Scardovia, Olsenella, and Parascardovia were enriched in D_ct. These findings describe ordered niche-associated differences, not longitudinal change within individuals.

### 5. Association-network differences between plaque and dentin niches

ASV-level association networks for D_sp and paired D_ct samples showed different topological structures (|ρ| >0.6; FDR <0.05). The D_sp network comprised 87 nodes and 112 edges, with low density (0.030) and higher modularity (0.42) (Fig. 3A). The D_ct network contained the same number of nodes but 168 edges, higher density (0.045), and lower modularity (0.31) (Fig. 3B). Several nodes annotated as TM7 lineages, Fretibacterium, Parvimonas, and Tannerella had higher connectivity in the D_ct network. Differential-association mapping also showed altered within- and between-module associations (Fig. 3C). These results indicate differences in correlation structure between the two niches; they do not demonstrate direct microbial interactions or causal rewiring.

**Figure 3.**
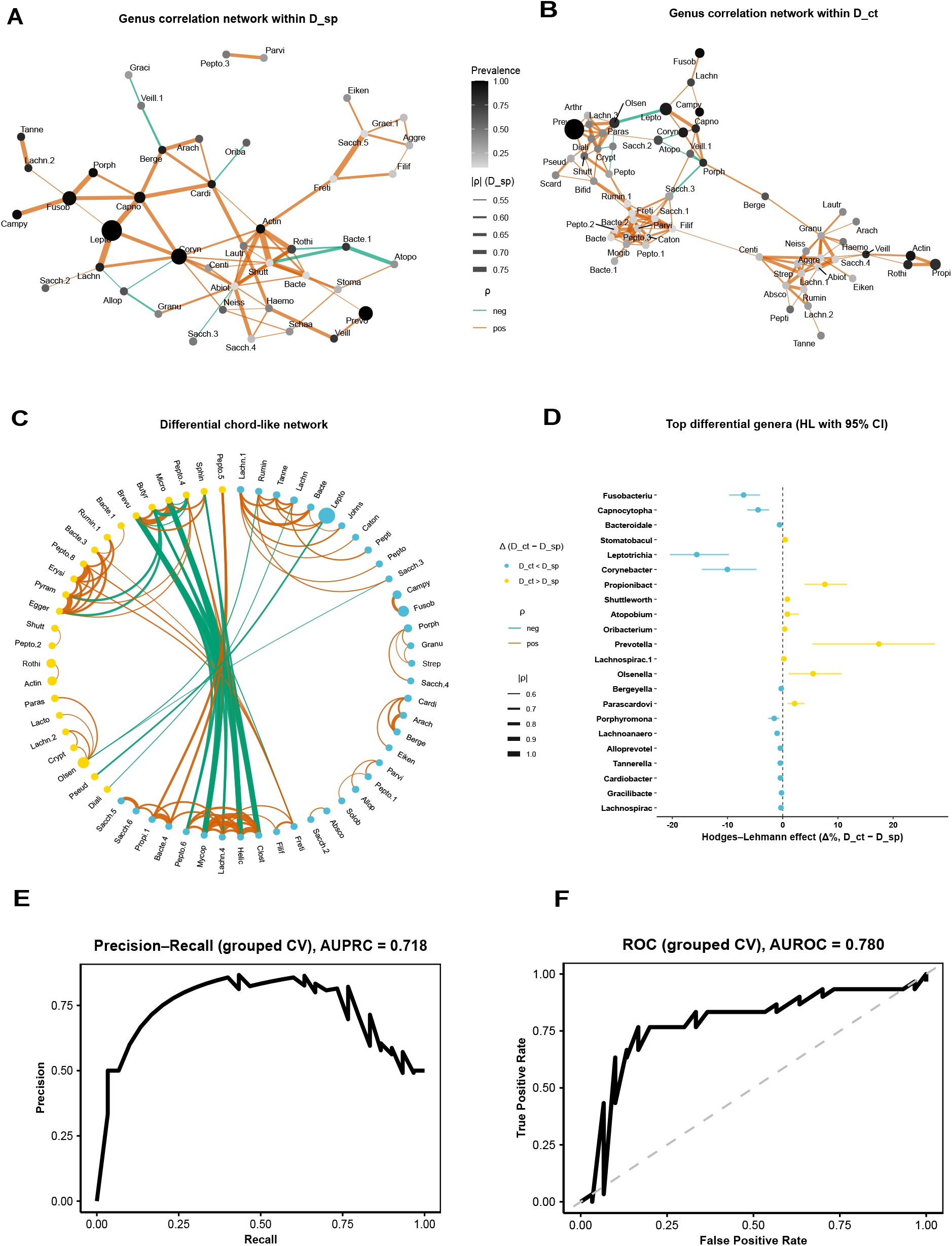
Association-network differences, differential taxa, and internal classifier performance across caries-associated oral niches. (A–B) ASV-level correlation networks in D_sp and D_ct, with nodes labelled by genus-level taxonomy. Node size represents prevalence; edge colour indicates correlation direction and edge width indicates association strength. (C) Differential chord-like network comparing D_ct and D_sp. (D) Top genera with differential abundance estimated by Hedges–Lehmann effects and 95% confidence intervals. (E–F) Patient-grouped cross-validation precision–recall and receiver-operating-characteristic curves for the sparse logistic-regression model. (G) Calibration plot for grouped cross-validation predictions. D_sp, caries-associated supragingival plaque; D_ct, carious dentin.

### 6. Network-associated taxa across niches

Paired differential analysis using CLR-transformed abundances identified 35 genera that differed significantly between D_ct and D_sp (FDR < 0.05), including 20 enriched in dentin lesions and 15 enriched in plaque (Fig. 3D). Effect sizes ranged from +15.2% for *Olsenella* to −12.4% for *Leptotrichia*, capturing both lesion-associated anaerobes and plaque-associated commensals.

Several lesion-enriched taxa, including *Olsenella, Parascardovia, Oribacterium*, and *Stomatobaculum*, exhibited concurrent increases in relative abundance and network centrality. In contrast, taxa such as *Fusobacterium, Tannerella*, TM7, *Leptotrichia*, and *Parvimonas* showed marked gains in network centrality despite higher relative abundance in plaque, indicating disproportionate structural roles within lesion-associated networks. Integrating abundance shifts with centrality measures, we defined a set of network-defined driver taxa comprising a lesion-enriched core (*Prevotella, Olsenella, Parascardovia, Oribacterium, Stomatobaculum*, Lachnospiraceae [G-7]) and a group of structurally prominent taxa (*Fusobacterium, Tannerella*, TM7, *Leptotrichia, Peptostreptococcus, Fretibacterium, Catonella*, and *Parvimonas*). Network features and ecological attributes of these taxa are summarized in Supplementary Table S1.

### 7. Sparse discrimination of plaque and dentin states

Using CLR-transformed genus-level profiles, we constructed a sparse logistic regression model to discriminate between D_ct and D_sp samples. The model was trained on paired samples and evaluated using patient-grouped 10-fold cross-validation. Selection of the λ_1_se solution yielded a parsimonious model incorporating three genera: *Olsenella* (positively associated with D_ct) and *Tannerella* and *Leptotrichia* (negatively associated with D_ct). The fitted model was defined as:

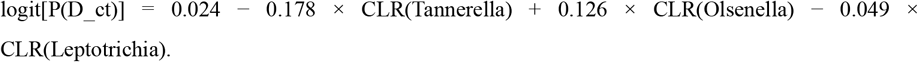

Model coefficients were consistent with both differential abundance and network centrality patterns, linking discriminatory features to ecological organization.

### 8. Internal cross-validation performance

In patient-grouped cross-validation, the model achieved an AUROC of 0.780 and an AUPRC of 0.718; the no-skill AUPRC was 0.333 (Fig. 3E–F). The calibration slope was 0.94 and the intercept was 0.03 (Fig. 3G). These estimates are internal validation results from a modest, single cohort and should not be interpreted as evidence of clinical diagnostic performance. Independent external validation is required.

## Discussion

This paired, cross-sectional study describes differences in oral microbial community composition and association-network structure across healthy supragingival plaque, caries-associated supragingival plaque, and carious dentin. The three sample types differed in diversity, taxonomic composition, and network topology [39–42]. Caries-associated plaque occupied an intermediate position in the ecological-ordering analysis, but this pattern should be interpreted as a cross-sectional niche relationship rather than a temporal sequence of disease progression.

The enrichment of Leptotrichia and Veillonella in caries-associated plaque, together with enrichment of Scardovia, Olsenella, and Parascardovia in carious dentin, is consistent with previously reported ecological differences between plaque and dentin lesions [42–46]. The observed diversity pattern likewise supports the view that oral niches differ in their physicochemical constraints. Longitudinal sampling will be required to determine whether individual communities follow the ordering observed here.

Association networks differed between plaque and dentin: the dentin network was denser and less modular than the plaque network. Such differences may reflect changes in co-variation patterns under distinct local conditions. However, compositional microbiome data and correlation-based approaches cannot establish direct microbial interactions; network results should therefore be interpreted as exploratory association patterns [32,33,49–51].

Integrating differential abundance with network centrality highlighted Olsenella, Parascardovia, Oribacterium, and Stomatobaculum as taxa associated with the dentin niche. Fusobacterium and Tannerella also had high centrality in the dentin network despite not being enriched in dentin. These observations identify candidate taxa for targeted validation, but centrality should not be equated with causal or keystone status without experimental evidence [52–54].

The three-genus logistic-regression model provided moderate internal discrimination between D_sp and D_ct samples. Its patient-grouped cross-validation design limits leakage from paired samples, but the model was trained and evaluated in one cohort and distinguishes sample niches rather than diagnosing caries in a clinical population. It should therefore be regarded as a proof-of-concept classifier. External validation in independent cohorts, with clinically relevant outcomes and prespecified thresholds, is required before any diagnostic or decision-support application.

This study has several limitations. 16S rRNA gene sequencing limits taxonomic resolution and does not measure function, which motivates metagenomic, metatranscriptomic, and metabolomic follow-up [55,56]. The healthy-control group was small and older than the caries group (median age, 35 versus 22 years), so residual age-related confounding cannot be excluded despite the non-significant age-group PERMANOVA result. The single-centre, cross-sectional design limits generalizability and causal inference. In addition, group dispersion differed in the beta-diversity analysis, and correlation networks should not be interpreted as direct interactions. Future longitudinal and multicentre studies should validate the observed niche associations and the candidate classifier.

In summary, paired sampling revealed distinct microbiome composition and association-network patterns in healthy plaque, caries-associated plaque, and carious dentin. Caries-associated plaque showed intermediate characteristics in several analyses. These findings provide a descriptive ecological framework and a set of candidate taxa for future mechanistic and externally validated studies.

## Conclusions

Healthy supragingival plaque, caries-associated supragingival plaque, and carious dentin showed distinct microbiome profiles in this paired, cross-sectional dataset. The intermediate position of caries-associated plaque in ecological-ordering analyses and the altered association-network structure in dentin provide hypotheses about niche-associated microbial organization. These results do not establish temporal progression, microbial causality, or clinical utility, but they identify candidate taxa and modelling features for validation in independent studies.

## Data Availability

Raw 16S rRNA gene amplicon sequencing data (V3–V4 region) have been deposited in the NCBI Sequence Read Archive (SRA) under BioProject accession number PRJNA1429009 and will be made publicly available upon publication. The submission includes 67 oral samples with corresponding BioSample and SRA run accessions.

## Acknowledgements

The authors thank all participants for their involvement in this study and acknowledge the staff of the Department of Stomatology, Xiangya Second Hospital of Central South University, for their assistance with participant recruitment and sample collection. We also thank BGI Genomics for technical support in sequencing.

## Conflict of Interest

The authors declare no conflict of interest.

### Author Contributions

Qingxiu Li and Lilan Zhao contributed equally to this work. Qingxiu Li performed data analysis, interpretation, and manuscript drafting. Lilan Zhao performed participant recruitment and sample collection. Zhenjun Li and Zhiwen Liu supervised the study and served as corresponding authors. All authors reviewed and approved the final manuscript.

---

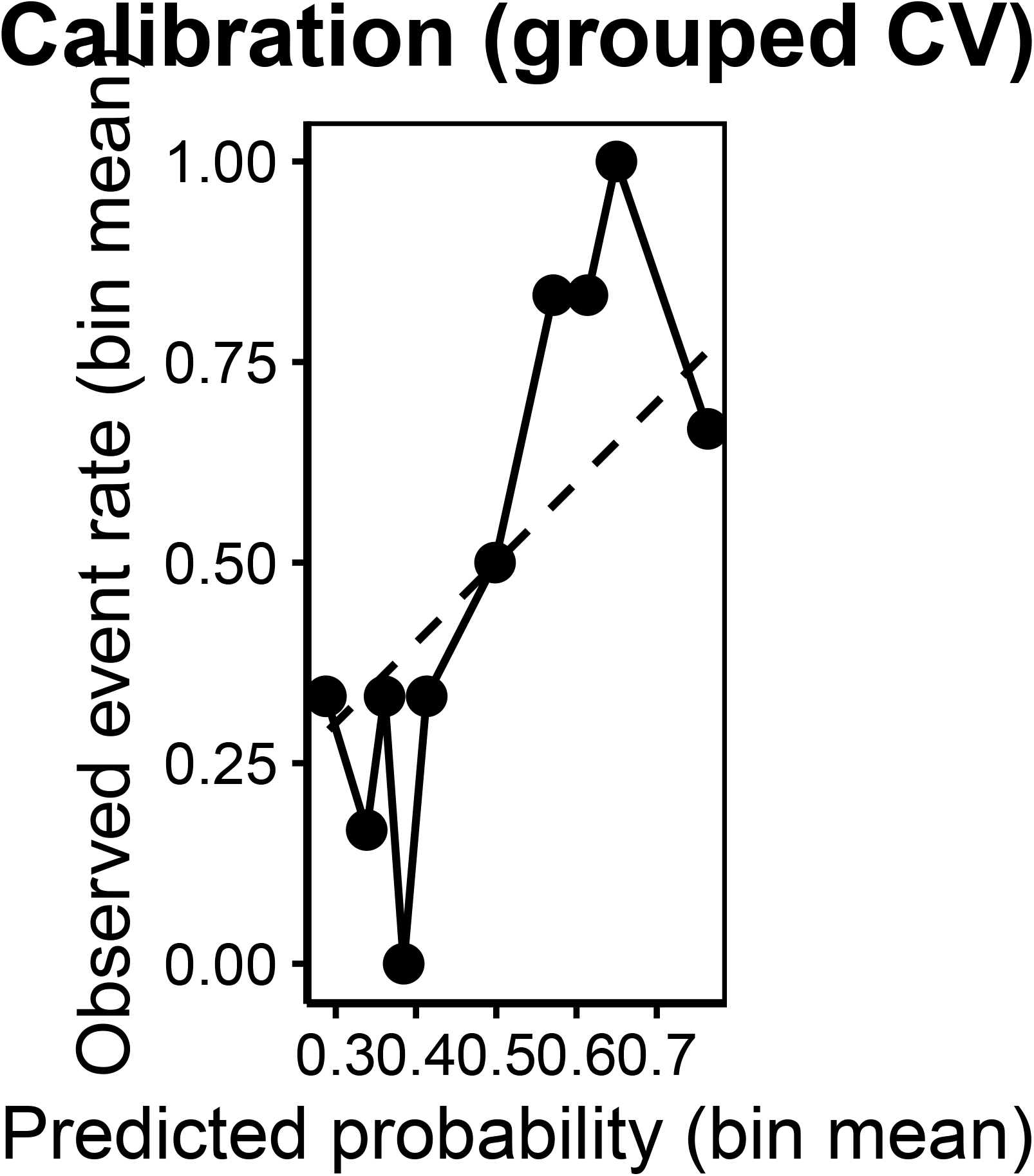

