## Supplementary Table S1 for "Ecological trajectories and microbial network reorganization across caries-associated oral niches"

### Supplementary Table S1. Driver taxa identified by integrated abundance and network analyses

| Taxon | Driver class | log2 fold change (D_ct vs D_sp) | FDR-adjusted q value | Abundance evidence | Degree centrality, D_sp | Degree centrality, D_ct | Change in degree centrality |
|---|---|---|---|---|---|---|---|
| Olsenella | Lesion-enriched core | 5.827 | 0.000 | D_ct-enriched | Not available | Not available | Not available |
| Parascardovia | Lesion-enriched core | 3.625 | 0.005 | D_ct-enriched | Not available | Not available | Not available |
| Stomatobaculum | Lesion-enriched core | 3.408 | 0.000 | D_ct-enriched | Not available | Not available | Not available |
| Lachnospiraceae [G-7] | Lesion-enriched core | 2.644 | 0.002 | D_ct-enriched | 0.059 | 0.161 | 0.102 |
| Oribacterium | Lesion-enriched core | 2.398 | 0.006 | D_ct-enriched | Not available | Not available | Not available |
| Prevotella | Lesion-enriched core | 1.113 | 0.003 | D_ct-enriched | Not available | Not available | Not available |
| Fretibacterium | Structural connector | 0.118 | 0.160 | No statistically significant abundance difference | 0.059 | 0.226 | 0.167 |
| TM7 [G-1] | Structural connector | Not available | Not available | Not available | 0.176 | 0.355 | 0.178 |
| Parvimonas | Structural connector | -0.868 | 0.781 | No statistically significant abundance difference | 0.088 | 0.226 | 0.138 |
| Fusobacterium | Structural connector | -1.632 | 0.001 | D_sp-enriched | Not available | Not available | Not available |
| Catonella | Structural connector | -2.032 | 0.040 | D_sp-enriched | 0.059 | 0.210 | 0.151 |
| Leptotrichia | Structural connector | -2.722 | 0.003 | D_sp-enriched | Not available | Not available | Not available |
| Peptostreptococcaceae [XI][G-7] | Structural connector | -2.939 | 0.003 | D_sp-enriched | 0.088 | 0.290 | 0.202 |
| Tannerella | Structural connector | -3.384 | 0.004 | D_sp-enriched | 0.029 | 0.129 | 0.100 |

Notes: D_ct, carious dentin; D_sp, caries-associated supragingival plaque. Log2 fold changes and q values were obtained from the paired D_ct-versus-D_sp differential-abundance analysis. Degree centrality values are reported only where the corresponding genus was retained in the network-centrality output; 'Not available' does not imply zero centrality. Lesion-enriched core taxa were selected from the integrated abundance/network panel. Structural connectors were selected from the same panel on the basis of their network role; their abundance evidence is reported separately.
